# Pressure and curvature control of contact inhibition in epithelia growing under spherical confinement

**DOI:** 10.1101/2021.06.17.448824

**Authors:** Ilaria Di Meglio, Anastasiya Trushko, Pau Guillamat, Carles Blanch-Mercader, Aurélien Roux

## Abstract

Morphogenesis requires spatiotemporal regulation of cell shape and proliferation, both regulated by biochemical and mechanical cues. In epithelia, this regulation is called contact inhibition, but disentangling biochemical from mechanical cues remains challenging. Here, we show that epithelia growing under confinement accumulate pressure that inhibits proliferation above a threshold value, which depends on the β-catenin pathway. Before inhibition of proliferation, cell aspect ratio abruptly increased upon reaching confluency. This shape transition occurred at low, constant pressure and was mainly controlled by cell density and contractility, correlating with YAP/TAZ pathway inhibition. In our system, epithelia spontaneously buckle: we observed that folding transiently reactivates both the YAP/TAZ pathway and cell proliferation. Altogether, our results support that different mechanical cues part of contact inhibition regulate cell proliferation through different mechanosensing pathways. Proliferation is regulated by sustained, tissue-level pressure through the β-catenin pathway, and by local curvature and pressure changes through the YAP/TAZ pathway.

## INTRODUCTION

The control of cell proliferation is essential for tissue formation and maintenance. In particular, which, where and when cells divide will contribute to final tissue shape, function and size. Once a tissue is formed, cells will continue to divide, mainly to compensate cell death and participate in tissue repair. Numerous physical cues, from extra-cellular matrix (ECM) stiffness to the compressive stress generated from tissue crowding, were shown to participate in the control of proliferation [1–5]. Cells must therefore be able to sense and respond to these cues by regulating the cell cycle to maintain the functional integrity of the tissue. Importantly, the inability of cells to sense or respond to these cues can lead to overgrowth, a hallmark of many diseases and development failures [6, 7]. In epithelia, contact inhibition englobes mechanisms by which a decrease in cell area, concomitant to increased cell-cell contacts, leads to cell proliferation arrest. Extensive evidence has supported the idea that an increase in cell-cell contacts is important but not sufficient for inhibition of proliferation [8, 9], yet we still lack a comprehensive picture of how mechanical cues control proliferation in tissues. This comes in part because mechanical cues are sensed through cell-cell adhesion protein complexes; one major mechano-sensing pathway is controlled through cell-cell adherens junctions, the α/β– catenin pathway, linking force and adhesion sensing at the molecular level. β-catenin, normally present at adherens junctions, shuttles to the nucleus after mechanical activation, binding T-cell factor/lymphoid enhancer factor (TCF/LEF) transcription factors to promote transcription of cell cycle genes [10]. Another signalling protein, the Hippo pathway effector Yap1 is involved in contact inhibition [9, 11] and is a general mechanosensing pathway [9, 12]. While β-catenin promotes progression into S phase, Yap1 nuclear translocation and binding to TEA DNA binding domain (TEAD) transcription factors promotes re-entry into mitosis [13]. Importantly, recent work showed that in response to mechanical stretch, Yap1 is transiently activated within one hour from stretch, whereas β-catenin activation occurs six hours from the stretch and is maintained for over sixteen hours [12].

Many studies propose that cell area may only represent a readout of the mechanical state of the cell [8, 14–17], which in turn regulates cell proliferation. In support of this, and as cell area is an easily quantifiable parameter, experimental studies using 2D cell culture models of epithelial tissues have posited that reduction in cell area in tissues growing under confinement regulates cell proliferation [4, 8, 9]. In this model, an increase in cell density would thus generate compressive stresses within the tissue that would inhibit cell proliferation. However, because of the difficulty to measure stresses within tissues, this assumption has not been fully tested. It is however established that increased mechanical tension promotes cell division [1, 5, 12, 15, 17], while increased mechanical compression generated by cell proliferation inhibits cell divisions [3, 18, 19]. But overall, how contact inhibition balances adhesive and compressive forces to regulate proliferation is not well understood.

Another process that is coupled to proliferation control is tissue curvature. Besides the well-known example of intestinal villi, where cells divide in concave crypts, and are extruded at the convex tip of villi, proliferation is coupled to curvature in many morphogenetic events. During formation of the *Drosophila* wing veins, areas with higher proliferation rates bulge out of the wing plane [20]. The confinement provided by surrounding less proliferative areas promote the buckling of the more proliferative veins through accumulation of compressive stresses. Overall, proliferation under confinement and tissue buckling are proposed to generate many of the curved shapes observed during tissue morphogenesis [21–23]. While measuring stresses in real tissues remains challenging [24–26], we have shown that compressive stresses generated within an epithelium growing under spherical confinement in vitro are sufficient to induce its buckling [27]. However, how the formation of folds regulates proliferation is still poorly studied. Interestingly, the Hippo pathway could be regulated by curvature, as YAP nuclear localization changes from concave to convex curvature in epithelium grown on wavy substrates in vitro [28].

Proliferation is the increase of cell number with time, and integrates new cells generated through cell division, and removal of old cells through cell death or extrusion. Here, we investigate whether compressive stress regulates epithelial cell division by assessing/measuring cell cycle progression under different pressure conditions. To this end, we employ an *in vitro* system based on the encapsulation and growth of an epithelial cell monolayer inside hollow hydrogel microspheres [18]. We then determine the relation between cell cycle progression and tissue pressure, which we measure by assessing the deformation of the elastic hydrogel shells. By using cells stably expressing FUCCI (Fluorescent Ubiquitination-based Cell Cycle Indicator), we show that cells stop dividing above 0.5-0.6 kPa. This threshold pressure value is dramatically reduced when interfering with the β-catenin pathway. Moreover, by monitoring cell aspect ratio, which is geometrically linked to cell density, we show a sharp increase in cell aspect ratio at the time when cells reach confluency. Interestingly, cell aspect ratio variations were found to be independent of pressure accumulation. Finally, we find that while YAP nuclear localization is inhibited when the cell aspect ratio changes, it is also transiently reactivated upon epithelium buckling, accounting for the transient reactivation of cell division observed in folds.

## RESULTS

### Cell encapsulation provides a system to study epithelial proliferation under spherical confinement

To investigate the link between mechanical forces and cell cycle dynamics, we monitored the growth of a model epithelial layer under spherical confinement. We encapsulated Madin-Darby Canine Kidney II (MDCK-II) cells inside hollow alginate microspheres, hereupon referred to as capsules, using a microfluidic device (Fig.1A) [29]. The microfluidic device consists of three co-centred channels that generate three-layered droplets: an outer layer composed of alginate (AL solution), a polysaccharide that undergoes gelation once the droplets fall into a CaCl_2_ solution; an inner layer with a suspension of MDCK cells and Matrigel (CS solution); an intermediate layer of sorbitol to avoid mixing between AL and CS (Fig. 1A and Methods). Upon capsule formation, the Matrigel forms a 3-4 μm thick layer on the inner surface of the capsules onto which cells spread and grow forming a closed monolayer [27, 29].

**Figure 1.**
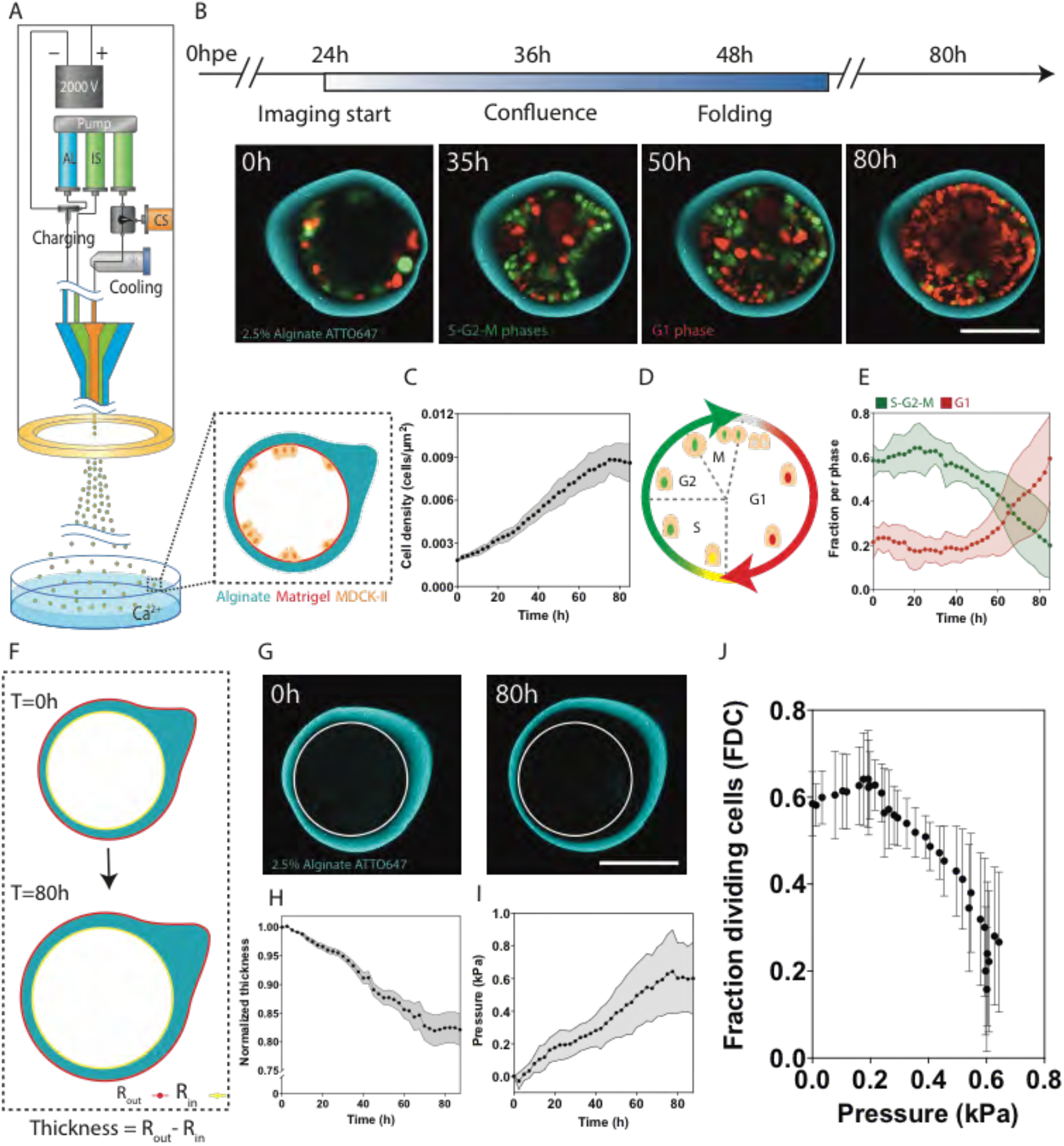
Cell cycle progression is regulated by pressure. (**A**) Schematic of the experimental setup (left) with schematic of capsule midplane at time of encapsulation (bottom right). **(B)** Timeline of growth of an MDCK monolayer inside capsules (top) and representative confocal equatorial plane images of MDCK FUCCI cells encapsulated in a 2.5% alginate capsule showing monolayer folding and capsule expansion (bottom). See also Movie S1. **(C)** Mean cell density (cells/μm^2^) over time for MDCK FUCCI cells encapsulated in 2.5% alginate capsules; 3 experiments, N=20 capsules. **(D)** FUCCI cell cycle indicator **(E)** Mean fraction of cells in G1 (red) and S-G2-M (green) phases over time inside 2.5% alginate capsules; 3 experiments, N=20 capsules. **(F)** Schematic of capsule expansion and wall thinning with inner (yellow) and outer (red) boundaries used to extract thickness change over time for pressure measurements. **(G)** Confocal equatorial plane image of alginate capsule at early and late time points showing capsule expansion over time; white circular contour shows the inner perimeter of the capsules at T=0h. See also Movie S2. **(H)** Mean normalized thickness of 2.5% alginate capsules over time; 3 experiments, N=20 capsules. **(I)** Mean pressure (kPa) over time for 2.5% alginate capsules. 3 experiments, N=20 capsules. **(J)** Mean fraction of dividing cells (FDC) as a function of mean pressure (kPa); 3 experiments, N=20 capsules. All error bars are SDs. Scale bars, 100 μm.

To assess the growth dynamics of MDCK cells, we used MDCK-II cells stably expressing the FUCCI reporter (Methods) in 2.5% alginate capsules starting at ~24h post-encapsulation (hpe) for a duration of 60-80h (times are given after imaging starts, AIS, Fig. 1B). Time-lapse experiments revealed that cells attach onto the capsule inner layer, grow to form a confluent monolayer that often undergoes folding and eventually fills the available space within the capsule (Fig. 1B, Movie S1). As we showed in our recent publication [27], compressive stress arising from cell proliferation of the confined epithelium drives folding through buckling. Even after folding, cell monolayers maintained epithelial polarity and organization (Fig. S1).

Cell density, which was obtained by quantifying cell number and inner capsule surface area (see methods), increased linearly for up to 60h AIS (Fig. 1C), supporting that neither confluence nor folding of the monolayer limited cell proliferation, which occurred 20-30h and 30-40h AIS, respectively. This linear growth rate of MDCK cells in capsules is in accordance with known growth dynamics of MDCK cells [30]. Cell density plateaued at 9±1 cells per 1000 μm^2^ 70-80h AIS, around 60h after confluence (Fig. 1C). This value is similar with the density steady state of MDCK monolayers grown on flat substrates [5]. However, cell density values for late stages of imaging correspond to folded monolayers and are not directly comparable to cell densities obtained on non-folded epithelia, as density is calculated from the capsule inner surface. Importantly, and as also proposed by numerous other studies [4, 8, 9], the observation that cell division rate decreases approx. 2-3 days after reaching confluence still supports the notion that cell-cell contact alone is not sufficient to inhibit proliferation. We next sought to quantitatively investigate the mechanical cues associated to confinement that could affect cell cycle dynamics.

### Cell cycle progression is halted past a threshold pressure

To probe how confinement affects the cell cycle, we used the FUCCI reporter (Methods) allowing us to monitor and quantify cell cycle dynamics; in G1 phase, nuclei are red, at the onset of S phase, nuclei become green and remain green throughout S-G2-M until the end of mitosis (Fig. 1D). Importantly, the accumulation of red nuclei can be caused by cell cycle arrest at the G1-S transition, or cell cycle exit into G0 [31]. We followed the evolution of cell cycle dynamics of encapsulated MDCK FUCCI cells over 80h (Fig.1C-E). For each capsule, we set t_0_ to the time at which cell density was in the range of 2 cells per 1000μm^2^, a value consistent with proliferating MDCK epithelia [4, 17]. Initially, cells actively cycle with most cells being green, thus in the S-G2-M phases (Fig.1B). As the monolayer grows and folds, we observe a shift to cells that are mostly red in G1-G0 phase, which suggests that confinement constrains cells to remain in G1-G0 phase (Fig. 1B).

We then quantified changes in cell cycle states over time, specifically the fraction of dividing cells (FDC), which is the ratio of S-G2-M cells over total cell number (Fig. 1E). For the initial 30h of monolayer growth, the FDC remains constant and much higher than the fraction of G1-G0 phase cells as most cells are actively cycling. FDC begins to drop at 40-50h, while G1-G0 cells fraction increases (Fig. 1E). The remaining cells at this point consist of cells just entering the S phase (yellow staining, Fig. S2). As confluence was reached at 18±6h (mean ± SD, N=25), the FDC remained constant even past confluence. Interestingly, the delay between time of confluence and the observed decrease in FDC was over 20h. Again, this confirms that confinement provided by capsules limits cell growth but that cell-cell contact alone is not sufficient for inhibiting proliferation. At approx. 50h, shortly before cell density plateaus (Fig. 1C), the FDC begins to decrease until it reaches almost undetectable levels at 80+ hours (Fig. 1E). The decrease in FDC is accompanied by a radial expansion of the capsules and a thinning of their walls suggesting that, as cells continue to divide, growth generates pressure that results in capsule deformation (Fig. 1F-G, Movie S2). In line with this, release of pressure by dissolution of alginate capsules with alginate lyase reactivated cell cycle progression showing that release of pressure restored cell growth (Fig. S3).

To assess the effect of pressure on cell cycle progression, we measured pressure accumulation from the growth-induced deformation of the elastic shell (methods and [18, 27]) (Fig. 1F-G). The change in capsule radius and wall thickness were computed from time-lapse images of the shell (Fig. 1G-H, Methods and [27]) and pressure was calculated over time from those changes (Fig. 1I and Methods). We observe that pressure accumulates linearly before reaching a plateau at maximum pressure of 0.6±0.2kPa at 75-80h (Fig. 1J). Notably, the drop in FDC corresponds to pressures between 0.4 and 0.6 kPa. Analysis of the FDC as a function of pressure reveals that as growth-induced pressure within capsules accumulates, the FDC decreases asymptotically towards the threshold pressure of 0.6kPa, above which cells no longer progress into the cell cycle (Fig. 1J). We conclude that confinement of an epithelial tissue generates compressive stress that impedes cell cycle progression past a threshold pressure of 0.6kPa.

### The effect of pressure on cell cycle progression is independent of capsule rigidity

To further show that pressure was the parameter regulating cell cycle, we encapsulated cells within softer shells to delay pressure accumulation, and quantify the evolution of cell cycle dynamics. Also, because substrate stiffness is known to regulate cell proliferation [9], we sought to assess whether modulating capsule rigidity affects cell cycle progression. By changing alginate percentage, the Young’s modulus (E) of the alginate capsule can be tuned [27]. Equatorial plane time-lapse images of 2% and 1.5% capsules illustrate the growth of the MDCK monolayer and the change in G1-G0 and S-G2-M phase cells over the course of 3-4 days (Fig. 2A&B). The measured Young’s moduli for 2.5%, 2% and 1.5% alginate capsules are respectively 19.5±0.7 kPa, 20.7±0.7 kPa and 11.5 ±0.4 kPa (Fig. 2C). Pressure accumulation clearly exhibited a delay for reduced stiffnesses (Fig. 2D): while 2.5% alginate capsules reached a maximum pressure of 0.6±0.2 kPa and 2% alginate capsules reached a maximum pressure at 0.5±0.2 kPa, 1.5% alginate capsules reach a pressure of only 0.2±0.1 kPa within the same time period of 85h. Interestingly, despite the delay in pressure accumulation of 1.5% alginate capsules, the rate of increase in cell density was similar for all percentages of alginate (Fig. 2E). In contrast, quantification of the different cell fractions revealed different cell cycle dynamics, which are thus dependent on capsule rigidity (Fig. 2F). For 2% alginate capsules, the FDC increased slightly up to 40h and began to decrease like in 2.5% alginate capsules but only reaching a minimum of 0.4, whereas the G1-G0 phase cells remain low and only increased slightly between 60-80h (Fig. 2F). In 1.5% alginate capsules, the initial FDC is lower compared to both 2.5% and 2% alginate capsules and remained between 40 and 50% FDC for the duration of the experiment (Fig. 2F). FDC exhibited a slow decrease as a function of increasing pressure but no drop for either 2% or 1.5% alginate capsules (Fig. 2G), as the pressure never reached the threshold pressure of 0.6kPa. However, in all cases, the curves of FDC with pressure followed the same path, albeit the lower value in 1.5% alginate (Fig. 2G). Overall, the fact that the drop of FDC is delayed in softer capsules, but that FDC as a function of pressure follows the same path for every stiffness confirms that pressure, and not capsule rigidity within the range used, regulates cell cycle progression.

**Figure 2.**
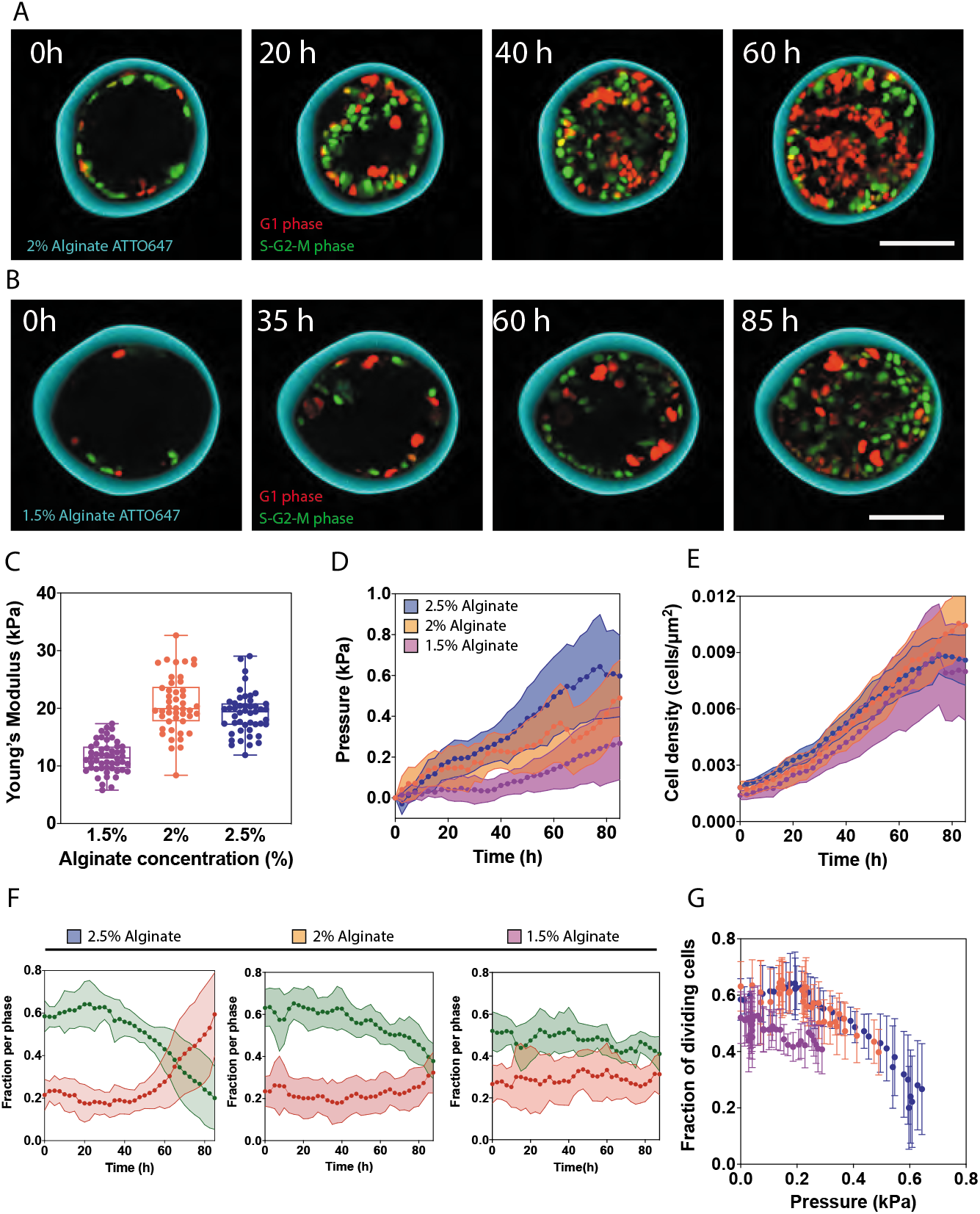
Pressure regulates MDCK cell cycle progression independently of substrate rigidity. **(A)** Representative confocal equatorial plane image of time-lapse for MDCK FUCCI cells encapsulated in 2% alginate capsules. **(B)** Representative confocal equatorial plane image of time-lapse for MDCK cells encapsulated in 1.5% alginate capsules. **(C)** Young’s modulus (kPa) for each alginate concentration measured using AFM. Respective Young moduli are: 1.5%, 11.5±0.4 kPa (N=52), 2%, 20.7±0.7 kPa (N=46) and 2.5%, 19.5±0.7 kPa (N=29). **(D)** Mean pressure (kPa) over time for 1.5%, 2% and 2.5% alginate capsules. **(E)** Mean MDCK FUCCI cell density (cells/μm^2^) over time for 1.5%, 2% and 2.5% alginate capsules. **(F)** Mean fraction of S-G2-M (green) and G1 phase (red) cells over time for 2.5%, 2% and 1.5% alginate capsules. **(G)** Mean fraction of dividing cells (FDC, S-G2-M cells) as a function of mean pressure (kPa) for 2.5%, 2% and 1.5% alginate capsules. Colors are the same as in panel D. For D-K, 1.5% alginate n=14, 2% alginate n=15, 2.5% alginate n=20. All error bars are SDs. Scale bars, 100 μm.

### Changes in cell aspect ratio do not depend on pressure

Previous studies [32, 33] have proposed that cell aspect ratio (AR) calculated as the ratio between cell height and cell width, would be a readout of pressure within the epithelium. Therefore, while cuboidal cells would feature AR≃1, compressed cells would be more columnar (AR>1) and stretched cells would be more squamous (AR<1). To test whether cells in capsules can accommodate lateral forces through AR variations, we measured AR over time and as a function of pressure, and for different capsule rigidities. To estimate the cell AR more accurately, we generated an MDCK-II stable cell line (methods) expressing both the cell membrane marker Myr-Palm-GFP and histone marker H2B-mCherry. Using this cell line, we manually measured the aspect ratio of 5-20 cells from fluorescence images of the midplane of 10-20 capsules per time point and per condition (Fig. 3A). Prior to monolayer confluence, cells are mostly squamous (Fig. 3A-B). As the monolayer reached confluence at around 4-5 cells/1000um^2^, cells became cuboidal, independently of rigidity (Fig. 3B). Past confluence, AR kept increasing indicating columnar cell morphologies for all capsule rigidities (Fig. 3B). However, the change of AR with time strongly depended on the rigidity of capsules, as cells in 2.5% alginate capsule changed their AR faster and more abruptly than for 2% and 1,5%, the latter being the slower, and the more continuous change of AR. We wondered whether the strong effect of capsule rigidity on the dynamics of AR could originate from rigidity itself, or from the lower accumulation of pressure in softer capsules.

**Figure 3.**
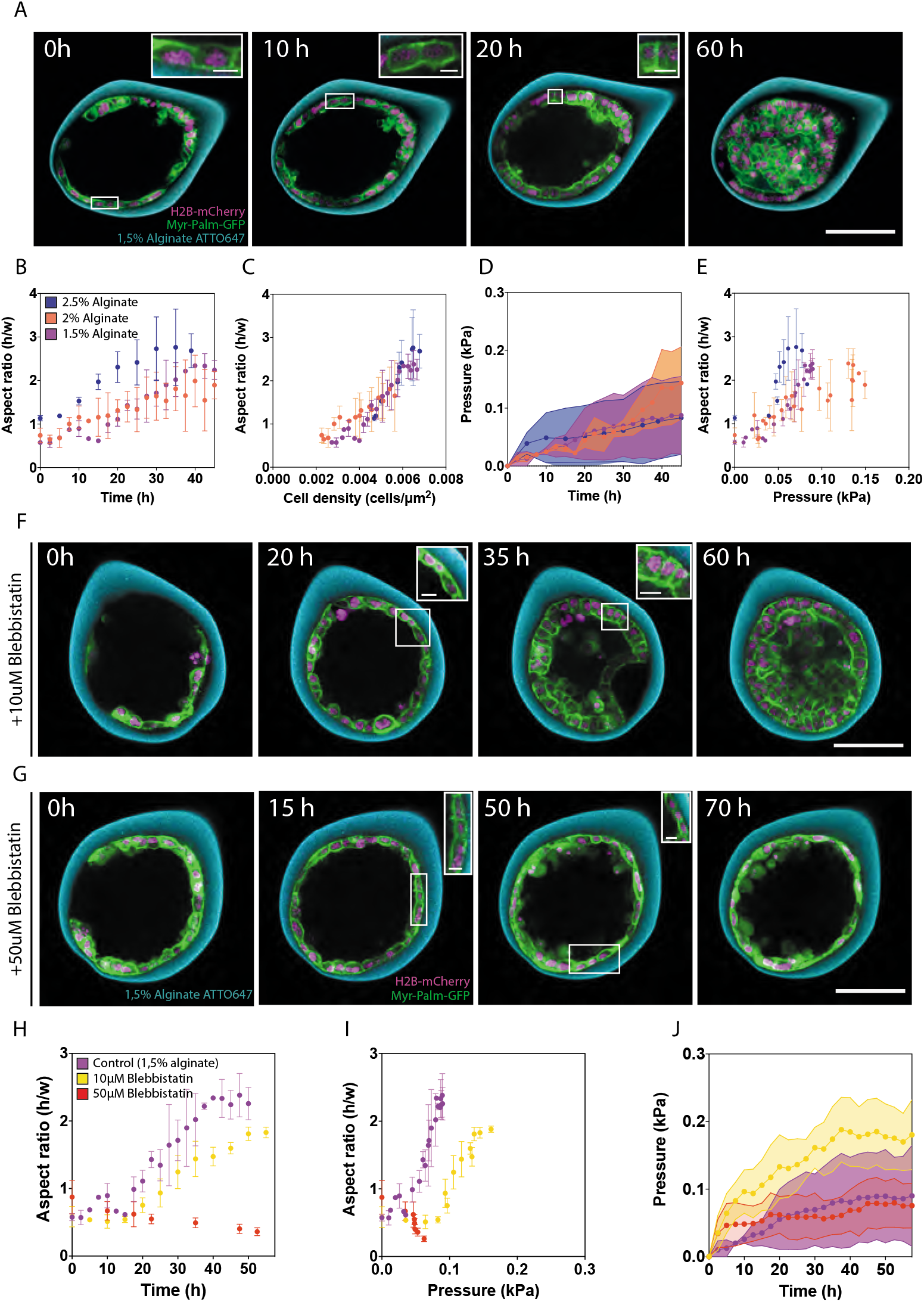
Changes in cell aspect ratio depend on substrate rigidity and cell contractility. **(A)** Representative confocal equatorial plane images of MDCK cells in 1.5% alginate capsules. **(B)** Mean aspect ratio (cell height/width) over time for different alginate percentages **(C)** Mean aspect ratio as a function of cell density (cells/μm^2^) for different alginate percentages **(D)** Mean pressure over time for encapsulated MDCK Myr-Palm-GFP H2B-mCherry cells. **(E)** Mean aspect ratio as a function of mean pressure for each alginate percentage **(F)** Representative confocal equatorial plane image of MDCK Myr-Palm-GFP H2B-mCherry cells encapsulated in 1.5% alginate capsules treated with 10μM Blebbistatin. See also Movie S3. **(G)** Representative confocal equatorial plane image of MDCK Myr-Palm-GFP H2B-mCherry cells encapsulated in 1.5% alginate capsules treated with 50μM Blebbistatin. See also Movie S4. **(H)** Mean aspect ratio over time for control and 10μM or 50μM Blebbistatin treated capsules. **(I)** Mean aspect ratio as a function of mean pressure (kPa) for control and Blebbistatin-treated MDCK Myr-Palm-GFP H2B-mCherry cells. **(J)** Mean pressure (kPa) over time for control and 10μM or 50μM Blebbistatin treated capsules. Colors in panels B-E corresponds to legend in B, and colors in panels H-J corresponds to legend in H. For B-E, 1.5% alginate n=14, 2% alginate n=18, 2.5% alginate n=19. For H-J, 10μM Blebbistatin n= 5, 50μM Blebbistatin n=10. All error bars are SDs. Scale bars, 100 μm; inset scale bars, 10 μm.

First, as expected from purely geometrical considerations, we noticed that the cell aspect ratio evolved linearly with cell density (Fig. 3C). Importantly, this linear dependence was independent from capsule rigidity, as the curves AR vs cell density perfectly collapsed for all three conditions of alginate percentage. We thus postulated that the difference in pressure accumulation in capsules of different rigidities may account for the difference of the cell AR dynamics (Fig. 3B). The Myr-Palm-GFP/H2B-mCherry cell line accumulated pressure rather slowly compared to the FUCCI and the H2B-mCherry cell lines used before [27](Fig. 3D), explaining why pressure accumulation was rather similar in all capsules, regardless of their alginate concentration (Fig. 3D). Consequently, when plotting AR vs pressure, dramatic differences in AR dynamics are observed with different stiffnesses: the larger the rigidity of the capsule is, the faster the AR changes (Fig. 3E). Also, AR reached higher values, and the transition to larger values of AR being more abrupt for stiffer capsules (Fig. 1E).

These results pointed to an internal mechanism of AR adaption independent of pressure, but dependent of capsule rigidity. Cell actomyosin contractility, which in MDCK cells is organized into an apical belt, is known to both participate in AR determination and to depend on substrate stiffness [1, 34, 35]. We thus wondered how cell contractility could influence cell AR dynamics during epithelium proliferation under confinement. For this, we reduced myosin-dependent contractility using Blebbistatin, and treated capsules with two concentrations, 10μM (Fig. 3F, Movie S3) and 50μM (Fig. 3G, Movie S4). At 10μM Blebbistatin, while the AR increased more progressively than non-treated cells, only reaching a moderate columnar shape (Fig. 3F). At 50μM, AR decreased with time, being more squamous at 70h than at 0h (Fig. 3G). Overall, reducing contractility strongly affected the dynamics and the value of the AR in Myr-Palm-GFP/H2B-mCherry (Fig. 3H&I), showing that cell contractility is a primary determinant of AR.

Interestingly, pressure accumulated faster in capsules treated with 10μM Blebbistatin than in non-treated capsules, suggesting that contractility limits pressure accumulation by promoting variation in the cell AR (Fig. 3J). However, when contractility is completely abolished with 50μM Blebbistatin, capsules accumulated almost no pressure, most probably because cell division was also affected at this drug concentration. Altogether, our results suggest that cell contractility is the primary control parameter of cell AR, and that pressure has little influence on AR. The dynamics of AR variations depend on capsule rigidity and independent of capsule pressure, consistent with cell contractility depending on substrate stiffness [9].

### Spatial distribution changes in cell cycle progression

We have previously shown that folding of epithelia growing in alginate capsules is due to buckling [27]. Curvature induction is associated with cell density and cell shape changes and could be associated with a local relaxation of the stresses, at least within the fold. We hypothesized these morphological changes may also affect cell cycle dynamics (see Fig. 1).

In order to assess how the temporal dynamics of the cell cycle is coupled to epithelial curvature, we sought to evaluate the spatial distribution of G1-G0 phase and S-G2-M phase MDCK FUCCI cells within folds (Fig. 4). Folded regions often featured mainly dividing cells (Fig. 4A). Quantifying numbers of resting and dividing cells in folds and outside the folds with time, we observe an increase in the fraction of dividing cells in folds, at time of folding (Fig. 4B, left), whereas regions outside the fold have a non-significant decrease of dividing cells at time of folding (Fig. 4B, right). As a consequence, at time of folding, the fraction of dividing cells within folds is significantly higher than in surrounding non-folded regions, a difference not observed when comparing non-folded regions of the same capsule (Fig. 4C). This is consistent with the notion that generation of curvature induces mechanical changes to the cell monolayer that triggers cell division transiently.

**Figure 4.**
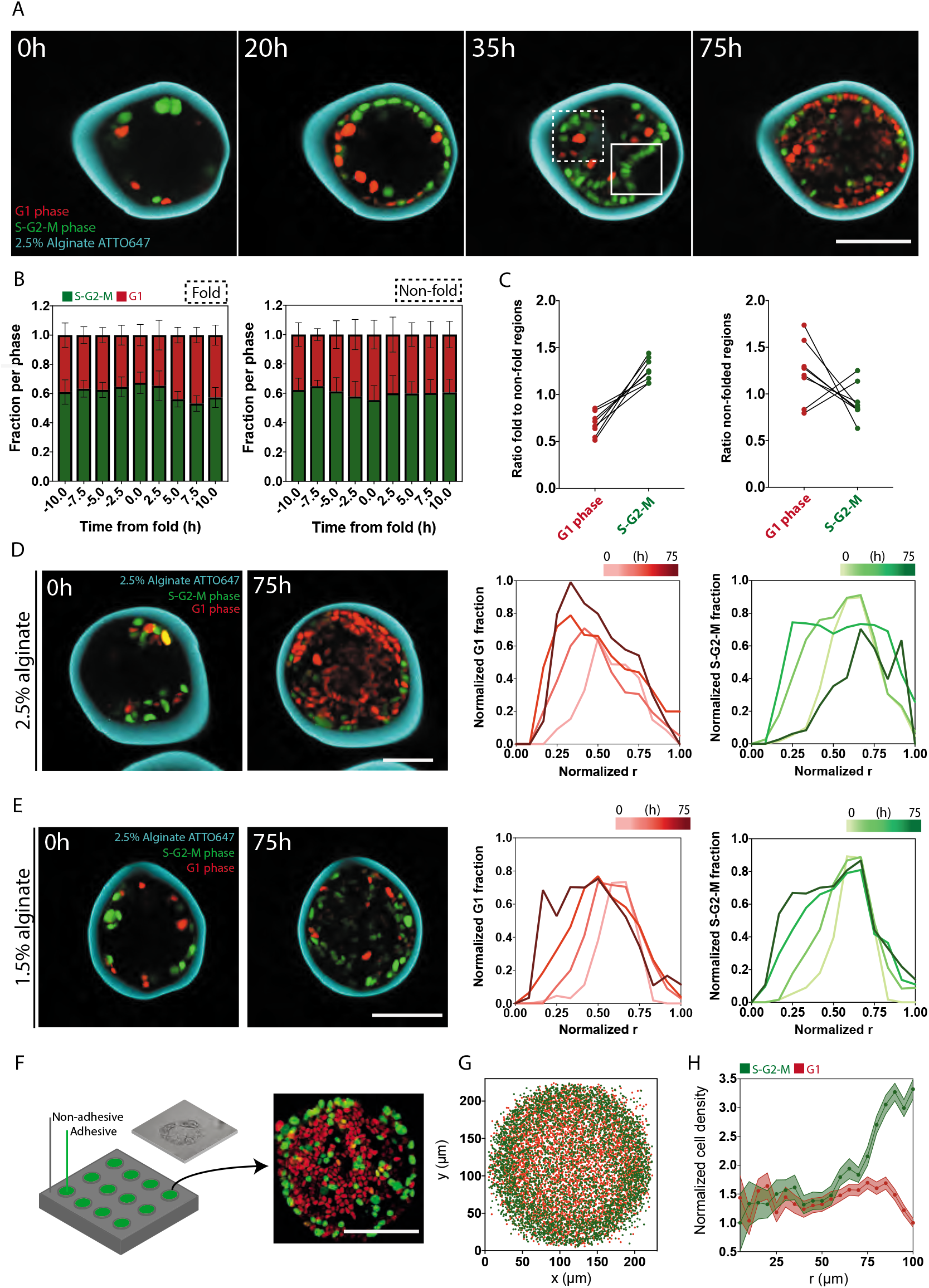
FDC distribution within folds and radial distribution of FDC. (**A)** Representative confocal equatorial plane image of time-lapse of MDCK FUCCI cells in 2.5% alginate capsules showing region of monolayer folding (solid inset) compared to non-folded region (dashed inset) used to compare the spatial distribution of FUCCI cells. **(B)** Fraction of cells in G1 or S-G2-M phases within fold (left) or outside of the fold (right) within 10h before and after time of fold (T=0h). **(C)** Ratio of G1-phase cells or S-G2-M phase cells in fold over non-folded region (left) compared to ratios in a non-folded region over another non-folded region (right). For B-C, N=8 folded and N=8 randomly chosen non-folded positions **(D)** Representative confocal equatorial plane image of MDCK FUCCI cells in 2.5% alginate capsules showing change in radial distribution of cell cycle phases at 0h and 75h and normalized fraction of cells in G1 (red) and S-G2-M (green) phases (normalized to max. fraction) as a function of radial distance from capsule centre (r: normalized to max radius) in time. 3 experiments, N=15 capsules. **(E)** Representative confocal equatorial plane image of MDCK FUCCI cells in 1.5% alginate capsules showing change in radial distribution of cell cycle phases at 0h and 75h and normalized fraction of cells in G1 (red) and S-G2-M (green) phases (normalized to max. fraction) as a function of radial distance from capsule centre (r: normalized to max radius) in time. 3 experiments, N=15 capsules. **(F)** Schematic of 2D adhesive circular patterns (left) with representative confocal maximum projection of MDCK FUCCI cells at over-confluence (about 4 days after seeding) on a circular pattern (right). **(G)** Position of G1-phase cells (red) and S-G2-M phase (green) cells on circular patterns at over-confluence. **(H)** Mean normalized cell density (to min. cell density) of G1 and S-G2-M phase cells as a function of the radius from the pattern centre (r=0 μm) to periphery (r=100 μm) of the pattern; N=78. All error bars are SDs. Scale bars, 100 μm.

To further study whether this transient activation of cell cycle during folding could impact the overall distribution of dividing cells, we followed the radial distribution of dividing and resting cells in time. We quantified the average radial distribution of dividing and resting cells by first normalizing cell positions by the final inner radius of the capsule, and averaged distributions over 15-20 capsules per time point (Fig 4D). Because capsules expand with time, cells are initially positioned at lower values than 1. In 2.5% alginate capsules, the peak of resting cell distribution progressively moved towards the centre of the capsule, whereas the peak of dividing cell distribution moved initially towards the centre of the capsule upon folding, and then moved back to the boundaries after folding (Fig. 4D). These results showed that the transient activation of cell cycle in folds impacts the overall distribution of cell divisions in the capsule.

At long times, the striking persistence of dividing cells at the 2.5% capsule boundary is reminiscent of previous observations in spheroids growing under osmotic pressure [3, 19]. It may be linked to a gradient of pressure within the forming tissue, lower pressure allowing for cell division at the periphery. To test this possibility, we used 1.5% alginate capsules, where pressure only reaches around 0.2kPa. In those capsules, both spatial distributions of dividing and resting cells slowly moved inwards, and no accumulation of dividing cells is seen at the periphery (Fig. 4E), in sharp contrast with the clear accumulation of dividing cells in the layer in contact with 2.5% capsules (compare Fig. 4D and 4E).

While maintenance of dividing cells at the periphery may be due to a pressure gradient, it could also be due to differences in adhesion, as cells at the periphery are adhering onto Matrigel, whereas cells in the core do not. Moreover, cells at the periphery are directly exposed to nutrients and growth factors diffusing through the alginate, while diffusion to cells in the core of capsules may be less efficient.

To test whether differences in cell adhesion or nutrient diffusion could generate maintain dividing cells at the periphery, we grew MDCK FUCCI cells on 2D adhesive patterns. In this case, cells are uniformly exposed to nutrients and growth factors, and all adhere to the substrate. In particular, cells were seeded on fibronectin-coated circular patterns with a diameter of 200μm (Fig. 4F, left, and Methods) and grown until over-confluent stages (around 72h) (Fig. 4F, right). Superimposing the nuclei signal of many patterns immediately showed that dividing cells are prominent at the periphery of patterns (Fig. 4G). Further quantification of the radial cell distributions revealed that resting cells concentrated in centre, whereas the periphery concentrated predominantly dividing cells (Fig. 4H). Thus, we concluded that neither detachment from the substrate, nor limited access to nutrients could be the cause of increased FDC at the capsule surface, since the same effect was observed on patterns. It also suggests that the lower pressure experienced by the cells at the boundary may as well favour cell division. The smaller nuclear area of cells at the centre of patterns compared to the periphery also suggests these are under increased compressive stress (Fig. S4). This is in line with previous studies, which revealed that cells grown on circular patterns establish patterns of tensional force that positively correlate with proliferation patterns; cells at the centre are under increased compressive stress and do not proliferate, whereas cells at the periphery do [9, 15].

### De-regulation of β-catenin activity alters how cell cycle dynamics respond to pressure

Finally, we sought to assess how pressure and curvature sensing may be transduced by known mechano-sensing pathways regulating the cell cycle. Two evident candidates are the YAP/TAZ pathway and the Wnt/β-catenin pathway effectors, Yap1 and β-catenin, respectively, both of which have emerged as important mediators of the mechanical regulation of growth [12, 36, 37]. The activity of the YAP/TAZ pathway can easily be followed by nucleo-cytoplasmic shuttling of Yap1 [38], while activity of the β-catenin pathway can be altered by phosphomimetic mutants and inhibiting peptides [36].

We first assessed the activation of the YAP/TAZ pathway in control conditions through nuclear localization of Yap1 over time. Encapsulated cells were fixed at pre-monolayer stage, at time of confluence and in late stages of folding and immunostained for Yap1 (Methods). Initially, nuclear localization of Yap1 was evident, but upon reaching confluence most nuclear localization disappeared, and stayed low on average for the rest of the process (Fig. 5A). These findings are consistent with the role of the YAP/TAZ pathway in establishing contact inhibition upon confluence and polarization of the epithelium [11, 39], but also suggest that Yap1 is not involved in the inhibition of cell cycle progression upon reaching the pressure threshold.

**Figure 5.**
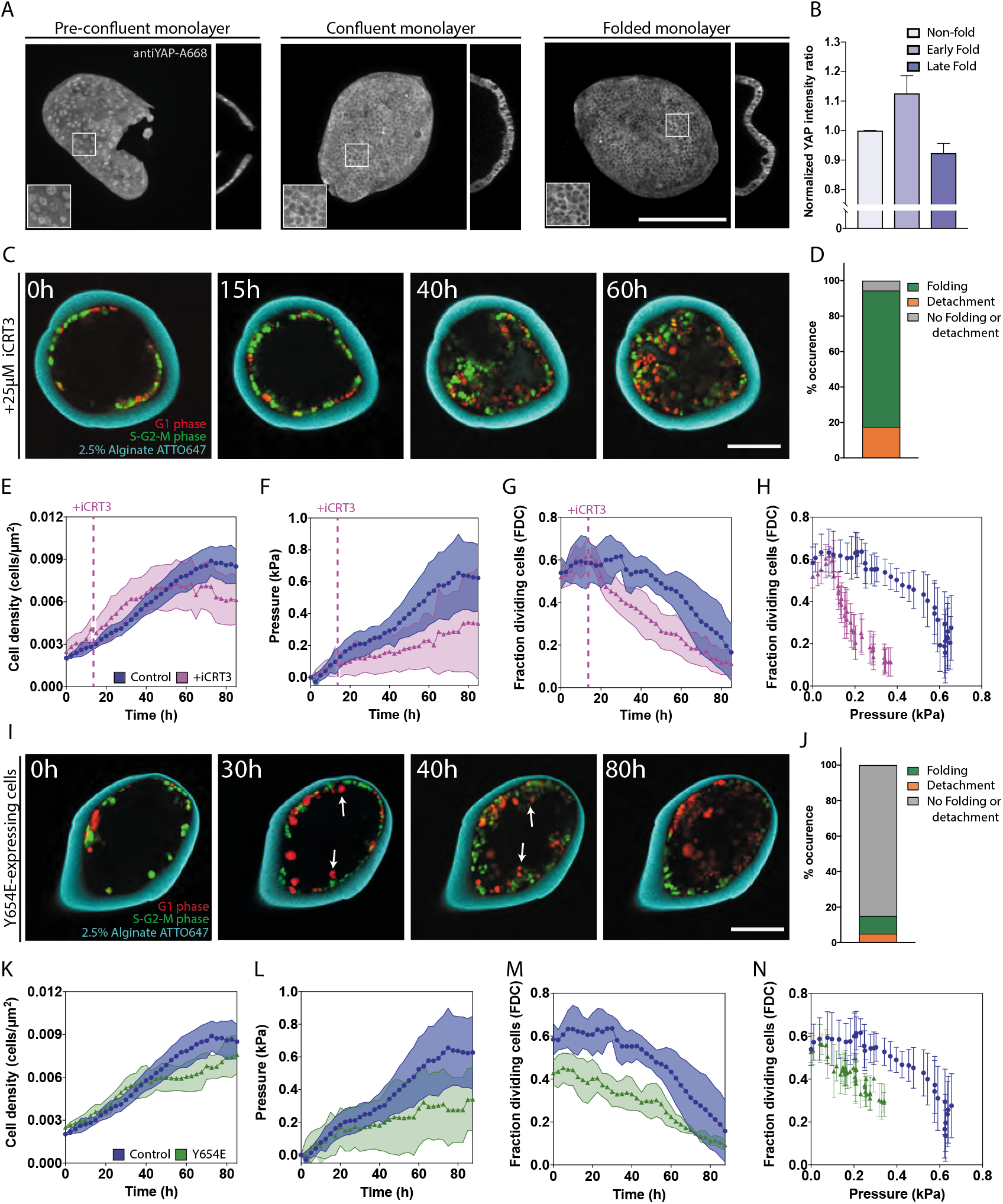
Inhibition and over-activation of β-catenin transcriptional activity alters pressure response. **(A)** Maximum z-projections and orthogonal view of fixed MDCK monolayers at preconfluence (left), at confluence (centre) and after folding (right) stage immunostained for YAP. (**B**) Normalized YAP intensity ratio (nuclear intensity of YAP normalized by mean nuclear YAP intensity in non-folded regions) non-folded (value 1) and early (approx. 24-30h AIS) and late (approx. 48h AIS) folded capsules. (**C)** Representative confocal equatorial plane image of time-lapse of MDCK FUCCI cells in 2.5% alginate capsules treated with iCRT3 at 15h and every 24h for the remainder of the experiment. **(D)** Percentage occurrence of folding, detachment or neither in capsules treated with iCRT3. **(E)** Mean cell density (cells/μm^2^) over time for control (circle, green) and iCRT3-treated (triangle, purple) MDCK cells within 2.5% alginate capsules. **(F)** Mean pressure (kPa) over time for control (circle, green) and iCRT3-treated (triangle, purple) MDCK FUCCI cells in 2.5% alginate capsules. **(G)** Mean fraction of cycling cells (FDC) for control (circle, green) and iCRT3-treated (triangle, purple) cells over time in 2.5% alginate capsules. **(H)** Mean fraction of cycling cells (FDC) for control (circle, green) and iCRT3-treated (triangle, purple) cells as a function of mean pressure (kPa) in 2.5% alginate capsules. (**I)** Representative confocal equatorial plane image of time-lapse of MDCK FUCCI cells expressing Y654E-beta-catenin-GFP in 2.5% alginate capsules. **(J)** Percentage occurrence of folding, detachment or neither in capsules Y654E-expressing FUCCI. **(K)** Mean cell density (cells/μm^2^) over time for control (circle, green) and Y654E-expressing FUCCI (triangle, orange) cells within 2.5% alginate capsules. **(L)** Mean pressure (kPa) over time for control (circle, green) and Y654E-expressing (triangle, orange) FUCCI cells in 2.5% alginate capsules. **(M)** Mean fraction of cycling cells (FDC) for control (circle, green) and Y654E-expressing (triangle, orange) FUCCI cells over time in 2.5% alginate capsules. **(N)** Mean fraction of cycling cells (FDC) for control (circle, green) and Y654E-expressing (triangle, orange) FUCCI cells as a function of mean pressure (kPa) in 2.5% alginate capsules. For B, n=3 per condition, error bars are SEM. For E-H, control n=20, iCRT3-treated n=12; error bars are SDs. For K-N, control n=20, Y654E-expressing FUCCI n=8; error bars are SDs. Scale bars, 100 μm.

Since Yap1 seemed involved in the early mechanosensing regulation of our system, we wondered if YAP could be involved in the transient re-activation of the cell cycle during folding. For this, fixed capsules in which folds were visible were analyzed, and the fluorescent amount of YAP in nuclei found in the folds compared to the ones found outside the fold (Fig. 5B and Fig. S5). We observed a small but significant increase of nuclear YAP1 in early folds, as compared to the non-folded parts, but older folds rather showed a small but significant decrease of nuclear YAP1 (Fig. 5B). These findings suggest that YAP1 is involved in triggering entry into mitosis of folding cells. These results also match the average movements of dividing cells in the radial direction during folding (Fig. 4D&E).

Given that the accumulation of pressure within capsules occurs on a timescale of days, and that the Yap1 regulation happened much earlier, we posited that β-catenin may be mediating the pressure-dependent cell cycle arrest at long times [12, 36]. To test this, we encapsulated MDCK FUCCI cells in 2.5% alginate capsules (Fig. 5C) and treated cells with iCRT3, an inhibitor of β-catenin binding to TCF. Treatment with iCRT3 did not slow down growth immediately compared to control monolayers (Fig. 5E), but growth reached the steady-state much earlier than in control conditions (40h instead of 70-80h). As a consequence, at steady-state, cell density was lower compared to non-treated capsules. This result suggested that inhibition of β-catenin binding to TCF may lower the threshold pressure at which cells stop dividing.

To further test this possibility, we quantified pressure accumulation (Fig. 5F) and the FDC with time (Fig. 5G). Consistent with proliferation reaching the steady-state faster, pressure accumulation was much slower in iCRT3-treated capsules (Fig. 5F). Accordingly, FDC initially increased like in control conditions, until iCRT3 treatment, after which the FDC showed a steeper but limited decrease (Fig. 5G). After this iCRT3-induced initial decrease, the FDC decreased continuously and faster than control conditions, consistently with the slower accumulation of pressure (Fig. 5F). When plotting FDC as a function of pressure, a dramatic difference is observed compared to non-treated cells, the lowest values of FDC not even reaching 0.4 kPa (Fig. 5H). This supports that the β-catenin transcriptional activity is required to set the threshold pressure.

We next sought to investigate whether altering the localization of β-catenin could also affect the response to pressure accumulation. We transiently expressed the phospho-mimetic mutant Y654E-β-catenin-GFP in MDCK FUCCI cells; phosphorylation of this residue is known to greatly reduce β-catenin affinity for E-cadherin [40], required for its translocation to the nucleus. We encapsulated Y654E-expressing cells within 2.5% alginate capsules and monitored monolayer growth and folding (Fig. 5I-J). Y654E-expressing cells had slower growth compared to control experiments, and did not reach the steady state within the 80h of the experiment (Fig. 5K. As consequence, pressure accumulation was slower than in controls (Fig. 5L). In Y654E-expressing cells, FDC was initially smaller than in control cells, and showed a constant decrease until 60h (Fig. 5M), explaining the overall slower growth and pressure accumulation. As a result, FDC decreased much faster with pressure than for the control (Fig. 5N), supporting that altering the capacity of β-catenin to shuttle between focal adhesions and the cytoplasm strongly interferes with the mechanical control of cell cycle. Altogether, these findings support that β-catenin transcriptional activity and shuttling activity are required for cell cycle regulation in response to pressure.

Interestingly, in iCRT3-treated cells and in Y654E-expressing cells, we observed a completely different morphogenesis outcome. While iCRT3-treated cells did fold almost normally upon reaching confluence (Fig. 5B), as seen in control conditions, Y654E-expressing cells did not fold at all, and stopped growing before folding (Fig. 5H). This suggests that in the case of Y654E-expressing cells, the threshold pressure at which cells stop growing is below the folding pressure, which we estimated to be 0.1kPa in previous works [27]. Another possibility is that Y654E-expressing cells adapt to pressure in a different way than folding, as we observed more cell extrusion in capsules with Y654E-expressing cells (Fig. 5I, arrows).

## Discussion

The spatiotemporal regulation of cell proliferation will determine the final shape and size of the tissue and it is crucial for both tissue morphogenesis and homeostasis. It has long been accepted that in addition to biochemical signals (i.e. growth factors), mechanical signals also contribute to the control of proliferation[3, 9, 12, 16, 36]. However, the mechanisms by which they do so remain unclear. The longstanding paradigm for proliferation control in multicellular tissues is a process termed contact inhibition of proliferation (CIP), which describes the tendency of cells to stop proliferating once the tissue reaches confluence [41, 42]. While tissue confluence eventually leads to practically undetectable levels of proliferation, CIP is a far more complex mechanism than was initially thought and it is now widely accepted that cell-cell contact alone is not sufficient to halt proliferation. A mechanism proposed extensively is that once cells establish cell-cell contacts, crowding could limit cell proliferation by imposing mechanical constraints on cell area.

While it is widely accepted that mechanical tension and compression will promote or hamper cell division [17, 43, 44], it is hard to quantitatively assess the link between mechanical stress and proliferation control in epithelial tissues. In this study, we show that cell cycle progression is regulated by compressive stresses arising from tissue growth under confinement. By measuring these stresses, we find that cell cycle progression halts past a threshold pressure in the range of 0.5-0.6kPa. This is surprising considering that previous studies on cells with no contact inhibition – such as cancer cells – showed that despite the pressure lowers proliferation, the latter is never completely inhibited [18, 19]. Our study establishes that the primary signal that controls proliferation during contact inhibition is sensing pressure increase. The response to pressure accumulation is directly altered when we either inhibit or over-activate the transcriptional activity of β-catenin, a transcriptional co-activator known to regulate cell cycle progression in response to mechanics [12, 36]. Also, the role of β-catenin, as well as its downstream effectors in sensing pressure and regulating cell cycle progression, emphasizes the importance of intact cell-cell contacts, where β-catenin mainly resides, providing with a molecular link between pressure sensing and contact inhibition.

In fact, Benham-Pyle *et al.* [36] previously showed that Wnt-dependent β-catenin activation and phosphorylation of β-catenin from mechanical stretch act synergistically to increase β-catenin-mediated transcription to levels required for mitosis. This could explain why in the case of over-activation of β-catenin using a phosphomimetic mutant, we do not observe continued or increased FDC and cell proliferation. Thus, given the consequences that a lack of cell cycle control will produce in a tissue, such as overgrowth in tumorigenic conditions, it is probable that other mechanisms, in parallel and in concert to the activity of β-catenin, come into play in response to high cell densities and increasing compressive stress. In line with this, previous work where MDCK monolayers are stretched, rather than compressed, revealed that iCRT3 treatment inhibits cell cycle [36]. This evidence further supports the notion that mechanical forces like stretching or compression regulate cell cycle progression in a β-catenin-dependent manner.

Another essential molecular pathway essential for contact inhibition is the YAP/TAZ pathway. In agreement with previous findings, we find that YAP nuclear localization changes upon reaching confluency, at a time when cells change their aspect ratio and polarization, but not their proliferation rate. This argues of a limited role of YAP/TAZ in triggering the proliferation arrest due to pressure, and an involvement of YAP/TAZ in rapid control of cell proliferation due to local changes of forces. β-catenin would then be involved in long-term and global sensing of mechanical forces. This possibility is supported by our finding that YAP/TAZ regulates early steps of contact inhibition, when cells reach confluency, whereas β-catenin regulates late stages of contact inhibition and pressure sensing.

In addition to the temporal regulation of cell cycle progression by accumulating pressure, we also find that cell cycle dynamics are spatially regulated within folds and across the tissue. Within epithelial folds that emerge through buckling, the FDC is transiently increasing. It is known that epithelium curvature correlates with different rates of cell proliferation, based on the observation of intestinal villi that stem cells are found in concave crypts, while cell death and extrusion is observed at the convex apex of villi [45–47]. In our system, however, it is hard to postulate whether the convex curvature of the fold promotes cell cycle progression – in opposition to the villi structure – or if it is rather the release of compressive stress that transiently re-activates cell cycle progression, but these two scenarios are also not mutually exclusive. The notion that lower compressive stress may maintain or promote cell cycle progression is also supported by the observation that dividing cells are only found at boundaries of the tissues in late stages of monolayer growth within capsules, an observation also made for cancer cells growing under confinement [3, 19]. However, this result made also be due to boundary effects, which imposes specific diffusion patterns of morphogens and hormones for cells found at the periphery of 2D circular micropatterns [48, 49] as well as 3D spheroids [50]. But a striking feature is the correlation we observed between this localised transient reactivation of the cell cycle, and the transient, localized re-activation of the YAP/TAZ pathway in forming folds. This further supports the notion that YAP/TAZ intervenes in sensing local, rapid changes of the mechanical environment of small groups of cells.

In contrast to cell cycle progression, cell aspect ratio does not depend on pressure accumulation but rather on subtle changes of substrate rigidity (5-20 kPa). Stiffer substrates allow cells to change their aspect ratio to higher values and faster compared to softer substrates. This change of aspect ratio strongly depends on acto-myosin contractility. A slight inhibition of contractility reduces the ability of cells to adapt to increased density by changing their aspect ratio but also results in increased pressure accumulation. This is suggestive of a mechanism by which cells may first reduce the accumulation of pressure by undergoing changes in cell aspect ratio by increasing contractility, this transition being under the control of YAP/TAZ. The balance between cell contractility (cell aspect ratio), adhesion to the substrate (which is known to be regulated by substrate stiffness) will determine the ability of the tissue to adapt to confinement. In the case where proliferation continues under confinement, this balance will determine whether folding or detachment of the epithelial tissue occurs [27]. When pressure becomes too high, cell aspect ratio cannot be further adapted, and proliferation is inhibited through the mechano-sensing activity of β-catenin.

Overall, our study proposes that epithelia growing under confinement can adapt to increasing pressure by initially changing their cells’ aspect ratio through contractility, then by folding if detachment from the substrate is allowed, and finally by inhibiting proliferation for pressure values above 0.5-0.6kPa. Our study establishes that cells with contact inhibition have a threshold pressure value above which they stop proliferating further validating recent theories [3].

## Acknowledgements

Authors thank Karsten Kruse for his useful insights into the project.

## Funding

AR acknowledges funding from Human Frontier Science Program Young Investigator GrantRGY0076/2009-C, the Swiss National Fund for Research Grants N°31003A_149975, N°31003A_173087, N°310030_200793 and N°CRSII5_189996. the European Research Council Consolidator Grant N° 311536. IDM and AR acknowledge funding from Secrétariat d’Etat à la Recherche et à l’Innovation grant agreement REF-1131-52107. I.D.M. and A.R acknowledge funding from the EU Horizon2020 Marie Sklodowska-Curie ITN “BIOPOL” (grant agreement No 641639).

## Author contributions

I.D.M. and A.R. designed the project; I.D.M. performed all experiments and image analyses, with exception of micropatterning experiments; A.T. helped with cell encapsulations, 4D confocal imaging and image analysis; P.G. performed 2D adhesive pattern experiments; C.B.-M. helped with image analysis; I.D.M. and A.R. analysed the results and wrote the paper, with editions from other co-authors.

## Declaration of interests

no competing interests stated.

## Notes

### Competing Interest Statement

The authors have declared no competing interest.

