## Supplementary Figures for "Pressure and curvature control of contact inhibition in epithelia growing under spherical confinement"

### Materials and Method

#### Cell culture and stable cell line generation

Madin-Darby Canine Kidney II (MDCK-II) (ECACC, Cat. No. 00062107) cells were maintained in DMEM (Invitrogen, cat. no. 10566016) supplemented with 10% (vol/vol) FBS (Thermo Fisher, Cat. No. 10270106), 1% (vol/vol) Penicillin-Streptomycin (Gibco BRL), and 1% (vol/vol) non-essential amino acids (NEAA) 100X (Invitrogen, ref. 11140050) in cell culture flasks (Falcon) at 37°C and 5% $CO_2$. Encapsulated MDCK monolayers were maintained under the same conditions as above and for live imaging transferred to MEM (Thermo Fisher, cat. no. 51200087) supplemented with 10% (vol/vol) FBS (Thermo Fisher, Cat. No. 10270106), 1% (vol/vol) Penicillin-Streptomycin (Gibco BRL), and 1% (vol/vol) non-essential amino acids (NEAA) 100X (Invitrogen, ref. 11140050) for image acquisition. The cell line MDCK FUCCI was a kind gift from the lab of Prof. Lars Hufnagel (EMBL Heidelberg, Germany). The cell line MDCK Myr- Palm-GFP was a kind gift from the lab of Dr. Mathieu Piel (Institut Curie, Paris, France), and was used to generate the cell line MDCK Myr-Palm-GFP H2B-mCherry. The plasmid H2B- mCherry, a gift from Robert Benezra (Addgene plasmid #20972), was inserted into the pLenti6.3/V5-DEST vector (containing C-terminal mCherry) using the Gateway cloning system. Lentiviral particles were generated in HEK293T cells using third generation lentiviral packaging vectors and MDCK Myr-Palm-GFP cells were infected with pLenti-H2B-mCherry. After infection, cell clones expressing both markers were sorted by Fluorescence Activated Cell Sorting (FACS) using a Beckman Coulter MoFlo Astrios, and monoclonal cells with unchanged morphology and sufficient expression level of the transgenes was selected. Cell lines were regularly tested negative for contamination with mycoplasma.

#### Microfluidic device fabrication

The microfluidic device (MD) was printed with the 3D printer EnvisionTEC Micro Hi-Res Plus, using the resin HTM140V2 (EnvisionTEC), with expected Z resolution of 25µm. The following printing parameters (set automatically based on the resin) were used: burn-in range thickness 400 µm, base plate of 300 µm, and exposure time 3000 ms. The printer light intensity is 225 mW/cm. The printed device was washed using ethanol and air dried using an air gun. A thin layer of PDMS (polydimethylsiloxane) at a ratio of 1:10 (curing agent: elastomer) was put on the cone of the chip with the help of a syringe needle and baked at 70°C for 30 minutes and subsequently baked using a UV chamber for 10 minutes. To ensure hydrophobicity and reduce the diameter of the device tip, a teflon tubing (PFA HP PLUS TUBING, 360 µm OD, 100 µm ID, IDEX) was used. The tubing was cut under a stereo binocular (Leica) using a scalpel to obtain a size of around 200-300 µm in length and glued on the tip of the microfluidic device with epoxyglue EA M-31CL (Loctite) and left to solidify for 1h at RT. To make the inlets, three 19-gauge stainless steel needles were cut into segments 1.5cm long and polished to avoid sharp edges. A small droplet of glue EA M-31CL was spread at the edge of the needles and they were inserted into the inlets of the devices, the glue was then left to solidify for 24 hours at RT.

#### Microfluidic device operation and cell encapsulations

The working principle of the microfluidic device is explained in detail in [27]. In brief, the system is composed of three syringes connected to a pump (Nemesys) for flow rate control, a Matrigel cooling part, the AL-charging part system and the microfluidic device. The MD consists of three coaxial cones inside which three different solutions are injected from the syringes. The outermost cone contains the alginate solution (AL), the intermediate cone contains 300mM sorbitol solution (IS) and the innermost cone contains cells/Matrigel/sorbitol solution (CS) in a ratio of 2:1:2 (v/v), with a cell number in the range of 5x10. The AL and IS solutions are loaded into two syringes controlled by the pumps for injection into the MD. The CS is injected into a cooling part to maintain Matrigel liquid, and this part is connected to a third syringe containing sorbitol that pushes out CS into the MD. The volume of Matrigel used was optimized to a total protein concentration of 0.2mg (optimized to ensure formation of a 3-4μm thick layer of Matrigel). The flow rates are set to 45 mL/h, 40 mL/h and 35 mL/h for AL, IS and CS, respectively, ensuring droplet formation upon exiting the MD. Once connected to the pumps the MD is positioned 50-60 cm above the petri dish with a 100mM $CaCl_2$ solution for collection of capsules. The alginate charging part and copper ring, both connected to a high voltage (2000V) generator, are used to improve capsule shape and monodispersity of size. The alginate charging part is a glass T connector that on opposite sides of the T has a high voltage wire (coming from the generator) and a tubing containing AL that flows down the T. The HV wire is coupled to a silver wire (OD 1 mm) that crosses the T such that it is in contact with the alginate and charges the solution, after which the charged AL then flows into the MD. The copper ring is held below the tip of the MD at a distance of about 1cm and centered with respect to the MD tip. The charged formed droplets passing through the copper ring under electrical tension get deflected as they cross the ring, creating a shower-like jet that prevents capsule merging. After 30 minutes in the calcium bath, capsules are washed and transferred to cell culture medium.

**Preparation of alginate solutions**

Labelling of 1% alginate solution with ATTO647N-amine (ATTO-TEC, ref. AD647N-95): 0.25 g Alginate (Protanal LF200FTS, FMS BioPolymer) was dissolved in 25mL 0.1M MES pH 6.0. Next, 5mg ATTO647N-amine dissolved in 200μL DMSO (anhydrous), was added into the tube and mixed, rotating for 30 min. Next, 21.5 mg sulfo-NHS (MW 217.14 g/mol, Fluka) dissolved in 200μL of 0.1 M MES pH 6.0 was added and let mix for 30min. Finally, 24mg EDC (MW 191.7 g/mol, Sigma) dissolved in 200ul of 0.1M MES pH 6.0 was added and let mix and react overnight at RT. The labeled alginate solution was transferred to a Slide-A-Lyzer$^{TM}$cassette 10K 12-20 mL capacity and let dialyze in miliQ for 2 hours, changing miliQ after the 2h and let to dialyze overnight. After dialysis, the labeled alginate was filtered (Acrodisc 25mm Syringe filter with 1um glass finer media, Pall, Life Science). The final concentration of the ATTO647N-labelled alginate was 0.55%, the solution was stored at 4°C. For preparation of 2.5%, 2%, or 1.5% ATTO647N-labeled alginate solutions, 0.27g, 0.165g, 0.22g of alginate powder (Protanal LF200FTS, FMS BioPolymer) is mixed with 10 mL milliQ water, respectively. Then, 1mL 0.5% ATTO647N alginate and 10 μl SDS 20% solution (Sigma, ref. 428018) are added and the mixture is left to rotate overnight at RT. Before use the solutions were spun down at 19,000 rpm for 30 min at 20°C, after which alginate was filtered with a glass fiber filter 1μm (Acrodisc 25mm Syringe filter with 1um glass finer media, Pall, Life Scince) before use.

**Micro-patterning**

Micropatterns were generated using the system PRIMO (Alvéole), mounted on an inverted microscope Nikon Eclipse Ti-2, via a UV-activated mPEG- scission reaction as described in [47]. After rinsing with PBS, the PEG-free PLL motifs were coated with Fibronectin (Sigma) at 50ug/mL for 5 minutes and then excess fibronectin was washed out. PBS was replaced with medium and MDCK cells were added. Samples were kept at 37°C and 5% $CO_2$. After 10-15 minutes, non-adherent MDCK FUCCI cells were washed out. Cells were left to grow to over-confluence for about 4-5 days before fixation.

**Immunofluorescence**

Cell monolayers inside capsules were fixed with 4% (v/v) paraformaldehyde (PFA) (Sigma, ref. F8775) in MEM (not PBS, to avoid dissolving alginate capsule) for 30 min at RT, then permeabilized and dissolved in 100 mM Glycine (Sigma, ref. G8898), 0.1% Triton X-100 (Applichemm, ref. A1388) and 1% Gelatin (Sigma, ref. G7765) in 1X PBS for 30 minutes at RT. Cells were then incubated with primary antibody diluted in 1X PBS with 1% Gelatin overnight at 4°C, then washed, and further incubated with secondary antibodies (1:1000) diluted in 1X PBS with 1% Gelatin for 1 hour at RT. Primary antibodies used for immunofluorescence staining were: rabbit anti-YAP (Cell Signalling, ref. 14074S). When necessary monolayers were also counterstained for f-actin and nuclei using Phalloidin488 (1:40, AlexaFluor488, Thermo Fisher, ref. A12379) and Hoechst 33342 (1:1000, Invitrogen, ref. H3570), respectively. Samples were rinsed 3X with 1X PBS.

**Inhibitor studies**

For contractility inhibition experiments, Blebbistatin (Sigma. ref. B0560) was dissolved in DMSO to a stock concentration of 17mM and used at a final concentration of either 10 μM or 50 μM (always freshly thawed from -20°C). Capsules were selected and pre-treated for 1 hour with Blebbistatin at 37°C and 5% CO2, after which they were embedded and imaged. For β-catenin-inhibition experiments, iCRT3 (Sigma-Aldrich, SML0211) was used to inhibit β-catenin/TCF interactions and transcriptional activity. iCRT3 was added 15 hours after imaging start (AIS) at a final concentration of 25μM and fresh inhibitor added every 24 hours until the end of the experiment.

**Image acquisition**

Capsules were first selected 24 hours post-encapsulation and embedded in 0.4% low-melting agarose (0.04 g in 10 mL) (Sigma, 49414) in a 35 mm glass-bottom dish (Mattek, Part No. P35G-1.0-14-C) to maintain them in the same position for several days of time-lapse imaging. The agarose was left to solidify for 15 minutes and 2-3mL of medium were added for imaging. For live fluorescence imaging, confocal images of samples were obtained using an inverted LSM780 NLO microscope (Carl Zeiss) using air objective 20x W Plan-APOCHROMAT 20x/1.0 DIC (UV) VIS-IR (Zeiss, 421452-9800). During imaging, capsules were maintained at 37°C with 5% $CO_2$. For each capsule, 3D confocal Z-stacks were acquired with 1μm intervals, and each capsule was imaged every 2.5-3 h for 25-40 cycles using definite focus (autofocus). For fluorescence imaging of fixed samples, images were acquired using the upright microscope LSM710 NLO 2-photon using an air objective 20X Plan-Apochromat 20x/0.8 M27 (Zeiss, 420650-9902-000).

**Capsule dissolution with Alginate Lyase**

For experiments assessing FUCCI response to capsule dissolution, FUCCI cell monolayers in capsules were embedded in 0.4% agarose for imaging in a 35 mm glass-bottom dish. The MatTek dish was manually modified by punching a hole on the side of the dish with a 19G hot needle to introduce teflon tubing into the dish sterilely for alginate lyase (Sigma-Aldrich, ref. A1603) injection. Once the MatTek dish was positioned onto the inverted LSM780 NLO microscope (Carl Zeiss) (section Imaging), 100 μL PBS alginate lyase solution (1 unit per 10 μL of PBS alginate lyase) was added both after 1min of imaging. The imaging was performed was in section Imaging. The imaging of capsules was performed with time intervals of 30 min for 8 hours and with definite focus (autofocus).

**Capsule pressure calculated from capsule deformations**

To measure pressure accumulation, confocal images of ATTO647-labelled alginate capsules were converted into binary black-white image masks using thresholding on Fiji (ImageJ). These masks were used to quantify pressure accumulation for encapsulated MDCK FUCCI cells as already described in [27].

**Image segmentation and processing**

The Imaris software (Bitplane) was used for 3D image visualization and nuclei segmentation from time-lapse confocal images. For quantification of MDCK FUCCI cell number and of S-G2-M and G1 phase cells, Imaris software is used to quantify intensity per channel (red and green individually). A custom-written MATLAB script was used to obtain cell number per phase and fraction of cells in either S-G2-M and G1 phase cells over time. For radial distribution analysis, a custom-written MATLAB script was used to define capsule centre and distance of nuclei from centre. For Yap1 analysis, nuclear segmentation was performed with Imaris software using Hoechst staining followed by pixel intensity calculation for each nucleus in the Yap1 channel. Using this method, cytoplasmic and plasma membrane staining were excluded, and the obtained nuclear pixel intensity of Yap1 was used to calculate a ratio of folds over non-folded regions.

**Statistical Analysis**

All statistical analyses for experimental data were performed in Origin (OriginLab Corp., Northampton, MA, USA). Statistics comparing two populations used unpaired t-test with Welch's correction. Results were considered statistically significant for p-values $\leq 0.05$. Error bars represent mean + SD (Standard Deviation) unless stated otherwise. Sample sizes are specified in figure legends.

### Supplementary Figures S1-S5

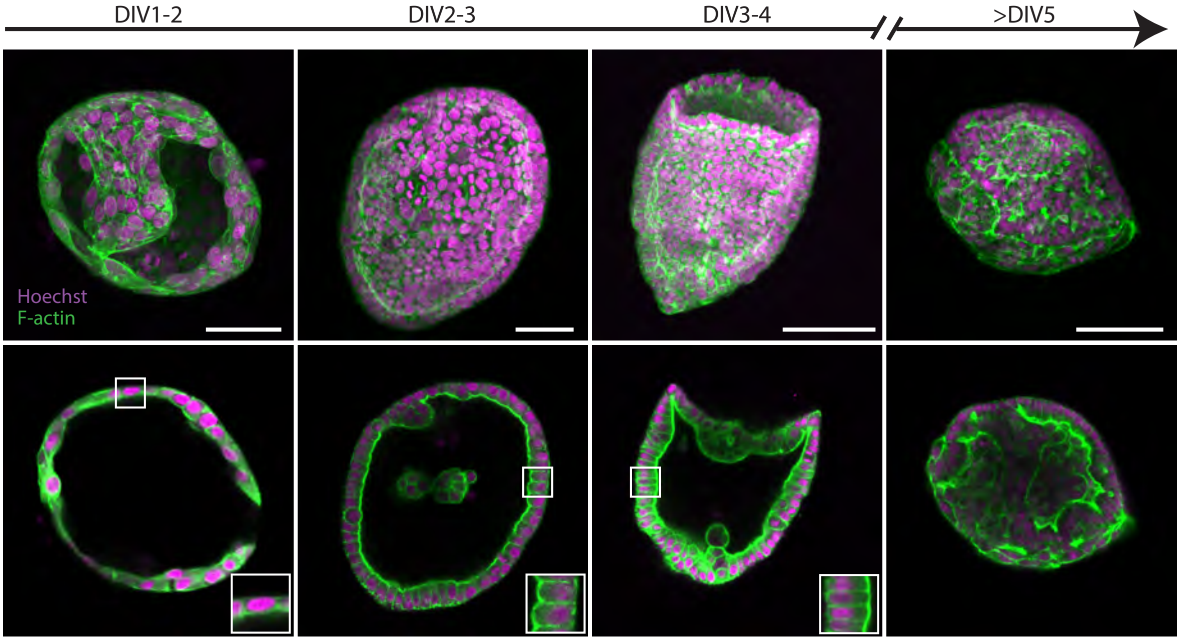


**Fig S1. MDCK monolayers maintain epithelial polarity and organization as they grow.** Related to Fig. 1. Confocal maximum projection (top) and corresponding equatorial plane images (bottom) of fixed and stained MDCK-II cells at different stages of monolayers growth inside capsules (not shown). Images are representative of at least 10 capsules per stage. at least 10 capsules per stage. DIV= days in vitro. Cells were stained for f-actin (green) using phalloidin-Alexa488 and nuclei (magenta) using Hoechst. Scale bars, 50µm

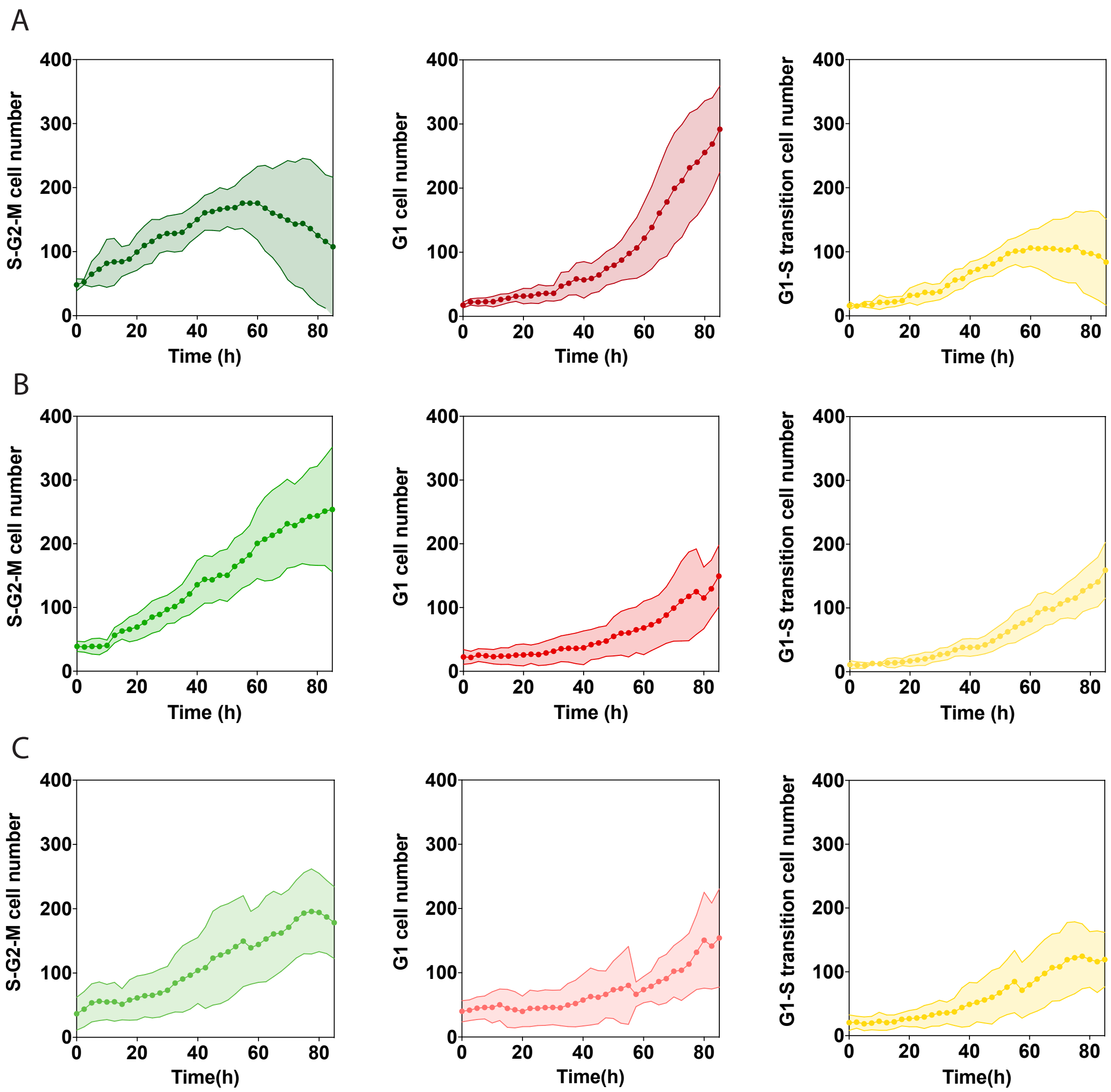


**Fig S2. Cell number per cell cycle phases for MDCK FUCCI cells in capsules of different alginate percentages.** Related to Fig. 2&3. Cell number in G1- phase (left), S-G2-M phase (centre) and G1-S transition (left) of MDCK FUCCI cells encapsulated in A 2.5% alginate capsules. B, 2% alginate capsules and C, 1.5% alginate capsules. Error bars are SDs.

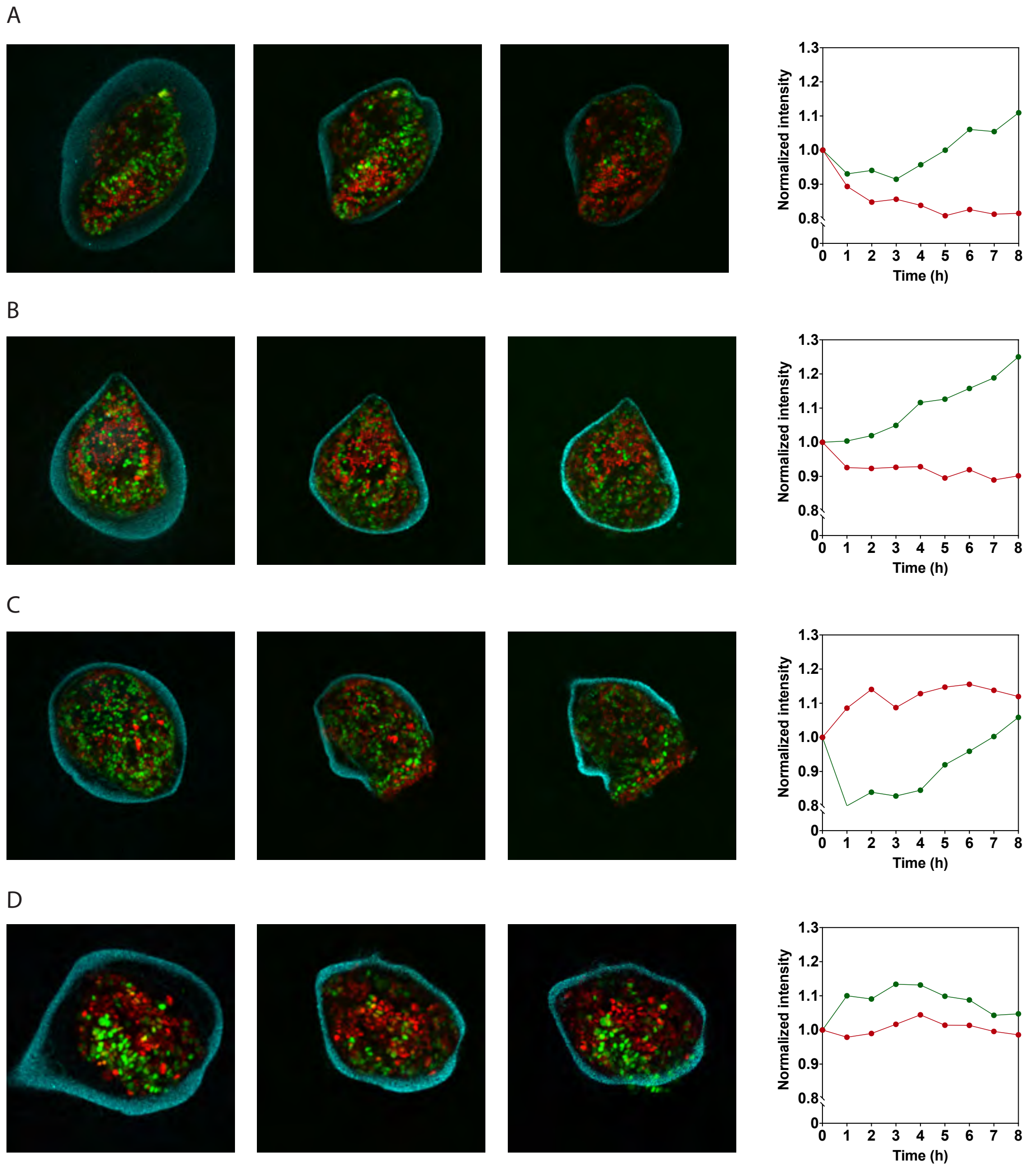


**Fig S3. Change in cell cycle fractions in over-confluent capsules dissolved with alginate lyase**. Related to Fig 2. Confocal maximum projection of MDCK FUCCI monolayers in response to capsule dissolution with alginate lyase. The intensity of maximum projection images was quantified to quantify change in G1 phase cells (red) and S-G2-M phase cells (green) from time of dissolution (0h) up to 8h after alginate lyase addition. Scale bars, 100 μm

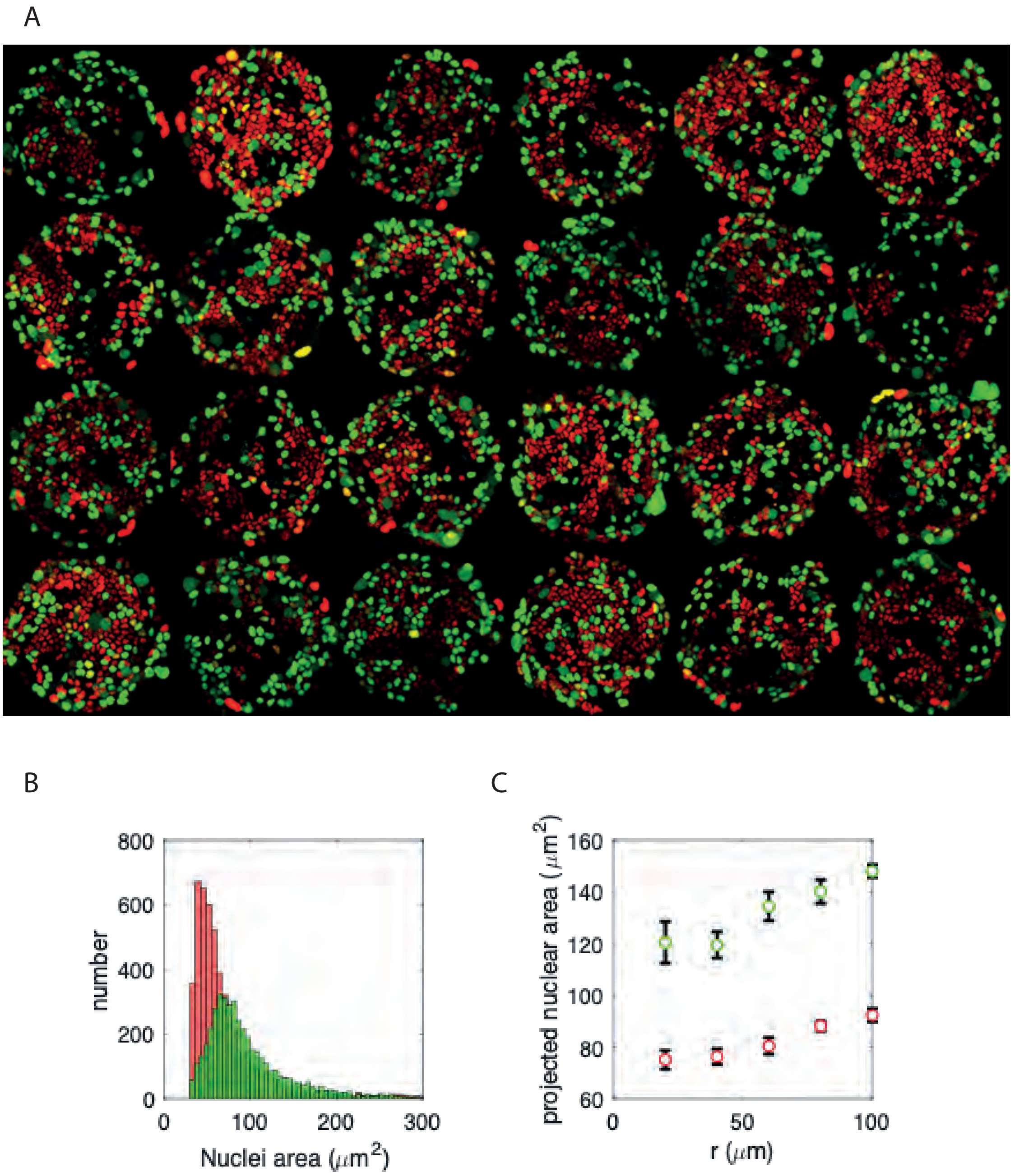


**Fig S4. Cell cycle dynamics of over-confluent MDCK FUCCI cells grown on 2D fibronectin-coated patterns.** Related to Fig. 3. A, montage of 24 representative maximum projection images of over-confluent MDCK FUCCI cells on spherical (200μm diameter) patterns. B, number of nuclei as a function of nuclei area (projected) for G1 cells (red) and S-G2-M phase cells (green). C, projected nuclear area for G1-phase cells (red) and S-G2-M cells (green) as a function of radius (0μm is centre of the pattern, 100μm is the border).

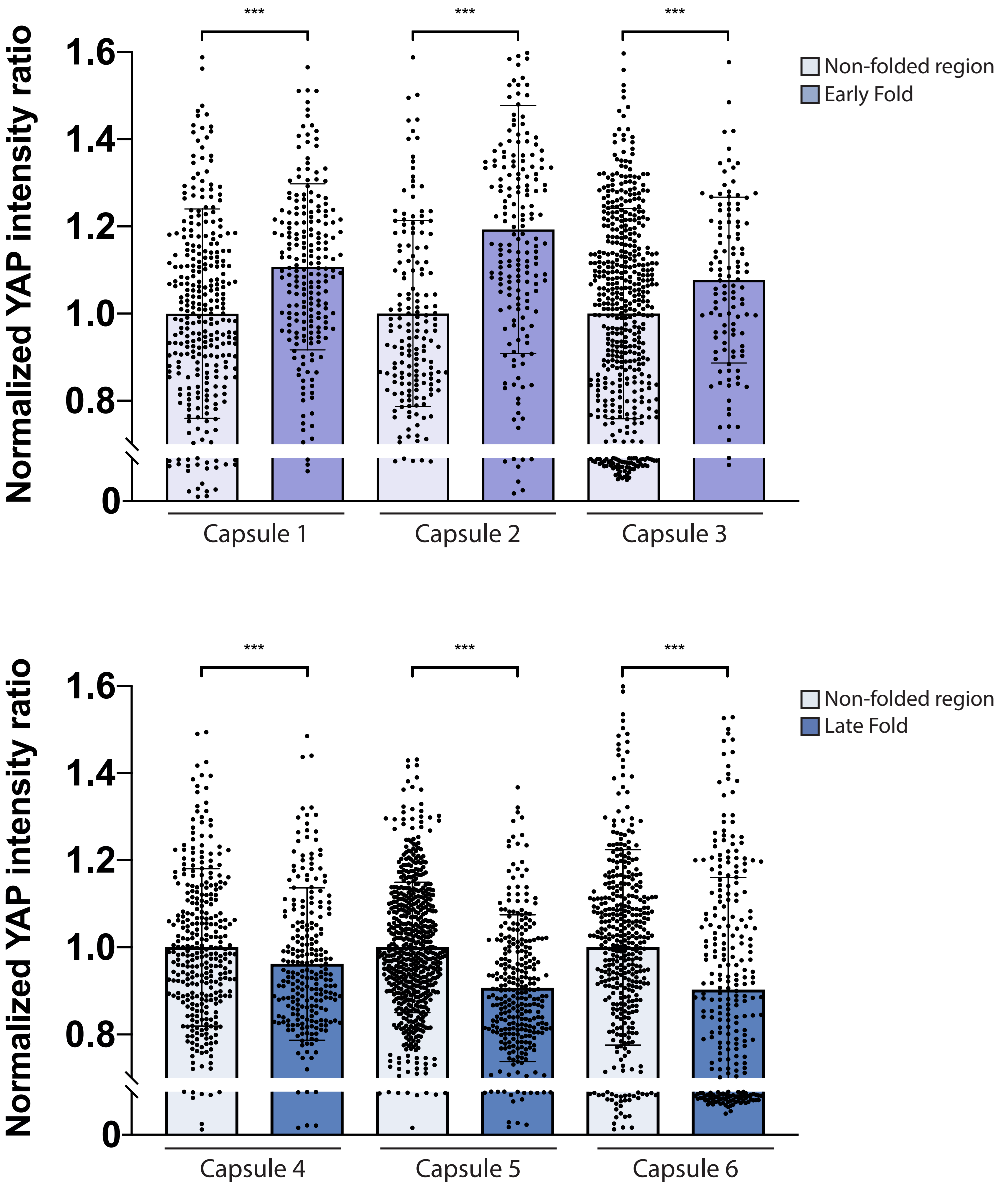


**Fig S5. Nuclear YAP increases transiently within folds.** Related to Fig. 5. YAP intensity ratio (intensity of nuclear YAP normalized to mean intensity of nuclear YAP in non-folded regions of the same capsule) for early folds vs. non-folded regions (top, capsules 1-3) and for late folds vs. non-folded regions (bottom, capsules 4-6).
